# Characterization of interstitial heterogeneity in the developing kidney

**DOI:** 10.1101/2020.03.02.973966

**Authors:** Alicia R. England, Christopher P. Chaney, Amrita Das, Mohita Patel, Alicia Malewsak, Daniel Armendariz, Gary Hon, Douglas Strand, Keri Drake, Thomas J. Carroll

**Affiliations:** Department of Molecular Biology, Center for Regenerative Science and Medicine, University of Texas Southwestern Medical Center, Dallas, TX, 75390, USA; Department of Internal Medicine, Division of Nephrology, University of Texas Southwestern Medical Center, Dallas, TX, 75390, USA; Hamon Center for Regenerative Science and Medicine, Dallas, TX, 75390, USA; Amgen, Inc., San Francisco, CA; Division of Pediatric Nephrology, University of Texas Southwestern Medical Center, Dallas, TX, 75390, USA; Department of Urology, UT Southwestern Medical Center, Dallas, TX 75390, USA; Laboratory of Regulatory Genomics, Cecil H. and Ida Green Center for Reproductive Biology Sciences, Division of Basic Reproductive Biology Research, Department of Obstetrics and Gynecology, University of Texas Southwestern Medical Center, Dallas, TX, 75390, USA

**Author notes:** Denotes equal contribution (Alicia R. England, Christopher P. Chaney, Amrita Das).

## Abstract

Kidney formation requires the coordinated growth of multiple cell types including the collecting ducts, nephrons, vasculature and interstitium. There has been a long-held belief that interactions between the progenitors of the collecting ducts and nephrons are primarily responsible for development of this organ. However, over the last several years, it has become increasingly clear that multiple aspects of kidney development require signaling from the interstitium. How the interstitium orchestrates these various roles is still poorly understood. We show that during development, the interstitium is a highly heterogeneous, patterned population of cells that occupies distinct positions correlated to the adjacent parenchyma. Our analysis indicates that the heterogeneity is not a mere reflection of different stages in a linear developmental trajectory but instead represents several novel differentiated cell states. Further, we find that beta-catenin has a cell autonomous role in the development of a medullary subset of the interstitium and that this non-autonomously affects the development of the adjacent epithelia. These findings suggest the intriguing possibility that the different interstitial subtypes may create microenvironments that play unique roles in development of the adjacent epithelia and endothelia.

**Graphical Abstract:** The developing interstitium is a highly heterogeneous, patterned population of cells that occupies distinct positions correlated to the adjacent parenchyma.

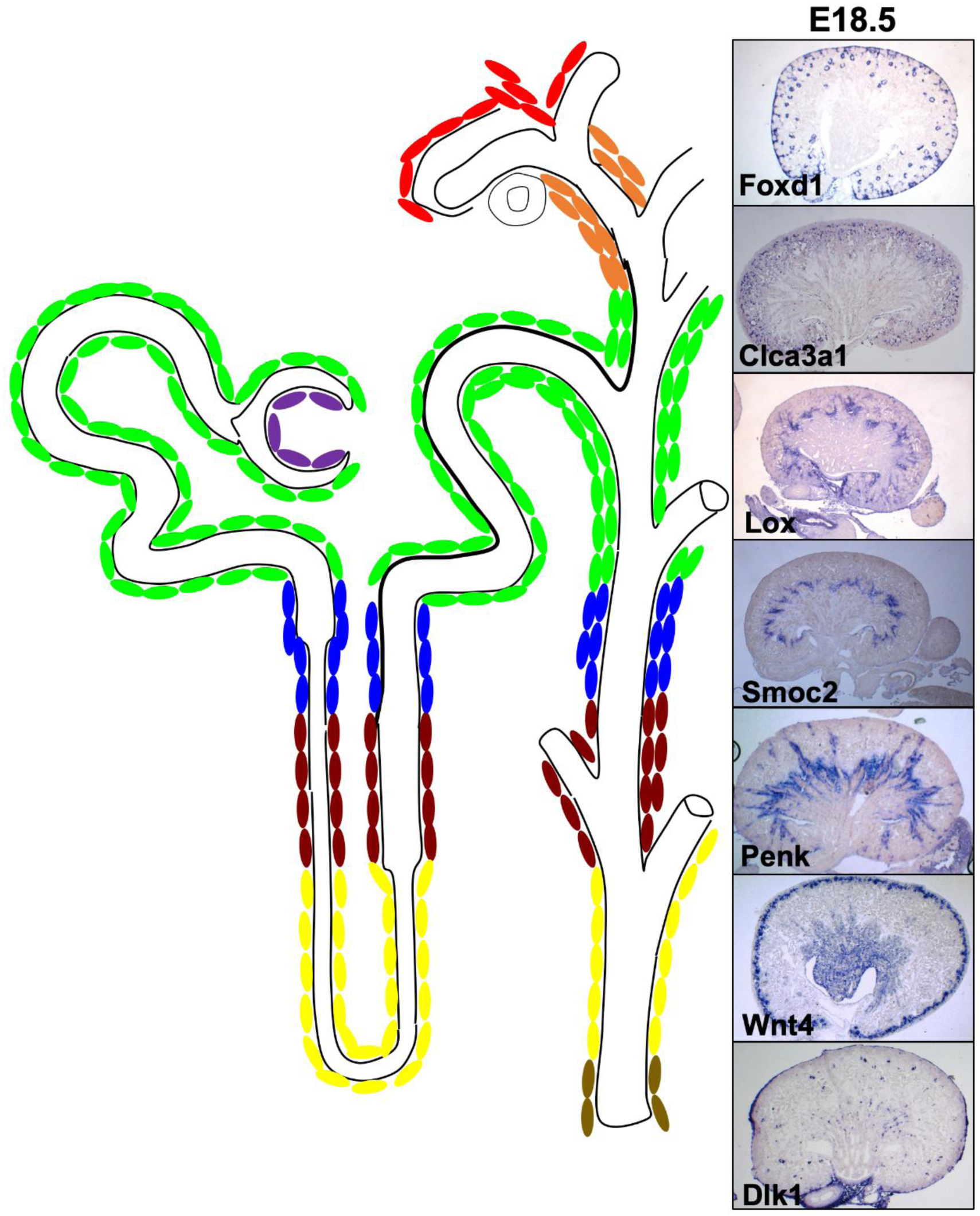

## INTRODUCTION

Development of the kidney relies on interactions between the metanephric mesenchyme (MM) and an epithelial structure known as the ureteric bud (UB) [1, 2]. The MM is a heterogeneous population of cells containing at least two cell type specific progenitor populations. A Six2/Cited1+ nephron progenitor cell (NPC) population is located within the MM directly adjacent to the tips of the UB [3, 4]. NPCs undergo mesenchymal-to-epithelial transition (MET) to ultimately form the nephron, the functional unit of the kidney, which is patterned into functionally distinct segments: the glomerulus, proximal tubule, loop of Henle, distal tubule and connecting segment [3, 4]. There is a second molecularly distinct progenitor population within the MM that surrounds the NPCs. These Forkhead box D1- (*Foxd1*, formerly known as *BF2*) expressing cells have been shown to give rise to a significant percentage of the interstitium and the renal capsule [5–7].

During development, the UB coordinates the proliferation and differentiation of nephron progenitors into the precursor of the nephron, the renal vesicle [8, 9]. Reciprocally, the MM (both the NPC and interstitial populations) induces the outgrowth and branching of the UB until the UB has formed an arborized epithelial network of tubules referred to as the collecting ducts [1, 10]. In addition to its role in regulating branching, the interstitium plays further roles in differentiation of the nephron progenitor population and patterning of the vasculature [5, 11–13]. Although frequently referred to in broad terms, the adult interstitial cell population includes renal fibroblasts and various smooth muscle cell types including vascular smooth muscle, pericytes, mesangial cells and the smooth muscle of the ureter and renal pelvis [6, 7]. Moreover, much of the endocrine function of the kidney, including the production of renin and erythropoietin, is performed by interstitial cells [14, 15].

Given that the interstitium has diverse roles in renal development, structure and function, it seems likely that this cell population is molecularly heterogeneous. However, extensive molecular characterization has been lacking. Here, we perform single cell RNA sequencing of Foxd1-derived interstitial cells from E18.5 mouse kidneys in order to define the heterogeneity and thus facilitate further inquiry into the development and function of these cells.

Our analysis revealed striking transcriptional heterogeneity in the renal interstitium, identifying 17 unique cellular clusters. Antisense mRNA *in situ* hybridization analysis demonstrated unexpected regionalization, uncovering at least 12 histologically similar but anatomically distinct domains along the cortical medullary axis. Importantly, comparison of mouse and human single cell data reveals that the interstitial heterogeneity is conserved. Analysis of transcription factor activity (regulons) showed cluster specificity, with the transcriptional regulator beta-catenin active within a medullary sub-population of stroma. Genetic ablation studies showed beta-catenin played a cell autonomous role in formation of this medullary interstitial region and a non-autonomous role in the development of the adjacent medullary epithelia. These findings stimulate multiple questions regarding the nature of interstitial-parenchymal crosstalk in development, physiology, homeostasis and disease of the embryonic and adult kidney.

## RESULTS

### Transcriptional analysis of the renal interstitium reveals molecular heterogeneity

Interstitial/stromal cells are present throughout the kidney extending from the cortex to the most medullary regions of the renal papillae and surrounding the ureter. Within the interstitium, three molecularly and anatomically distinct regions of interstitial cells have previously been defined. These populations were annotated as nephrogenic, cortical and medullary interstitium based on their unique anatomical positions [16]. However, more detailed examination of kidneys stained with various antibodies to proteins expressed in interstitial cells suggests that the degree of heterogeneity may be under-estimated [7]. For example, Foxd1 is expressed in a small subset of the interstitial cells cortical and lateral to the Six2-positive cap mesenchyme. Expression ceases adjacent to the renal vesicles (Figure 1a-b). In comparison, Tenascin-C is expressed in a subset of Foxd1 expressing cells lateral to the Six2-positive cap mesenchyme but is not expressed in the cells cortical to the cap. Tenascin-C expression extends into the cells adjacent to the proximal tubule (Figure 1c-d). Slug expression appears to be largely, if not completely, non-overlapping with Foxd1, marking a subset of interstitial cells just medial to the cap mesenchyme and extending medially to the level of the proximal tubules (Figure 1 e-f). Acta2 is detectable in the interstitial cells adjacent to the proximal tubules but is not detectable in the medullary interstitium of the renal papillae (Figure 1g-h). CDKN1c is expressed in the majority, if not all, interstitial cells of the renal papilla. Lef1 is expressed in interstitial cells extending from just medullary to the renal vesicles through the entire papillae (Figure 1i-j). However, in contrast to CDKN1c, Lef1 appears to only be expressed in a single layer of interstitial cells that lie directly adjacent to the collecting ducts (Figure 1k-l). Tbx18 is only expressed in interstitial cells surrounding the renal pelvis and ureter (Figure 1m). Interestingly, of all the proteins discussed, only Acta2 is expressed in the smooth muscle surrounding the vasculature and mesangial cells, both of which are Foxd1-derived interstitial cell types. Based on the spatial differences in expression of the proteins described above and recent studies, one would predict that there are may be as many as 10 distinct interstitial cell types within the kidney.

**Figure 1:**
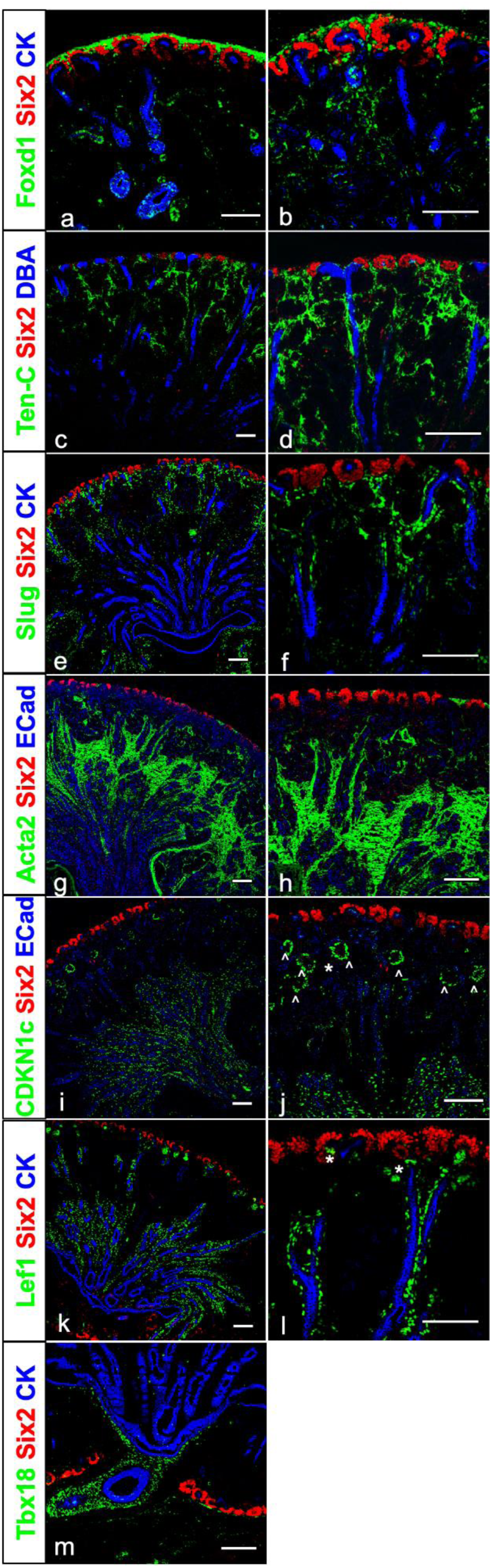
Unexpected heterogeneity within the renal interstitium revealed through antibody staining and single cell RNA sequencing. Wildtype E18.5 kidneys stained for Six2 and CK (red, blue respectively in a-m), and Foxd1 (green in a-b), Tenasin C (green in c-d), Slug (green in e-f), Acta2 (green in g-h), CDKN1c (green in i-j), Lef1 (green in k-l), Tbx18 (green in m). Scale bar 100um.

To gain a more complete understanding of interstitial heterogeneity within the developing kidney, we performed single cell RNA sequencing (scRNA-seq) on dissociated E18.5 wild type mouse kidneys. Although previous single cell analyses have been performed on both adult and embryonic kidneys [17–20], the interstitium represents a relatively small percentage of the total number of cells and thus has been under-represented. To enrich for interstitium, we purified cells via fluorescence activated cell sorting (FACS) from Foxd1Cre;Rosa-Tomato kidneys as Foxd1-positive cells have been shown to give rise to the majority of the renal interstitium [7, 21]. Further, because recent studies suggest that a sub-population of the interstitium (in particular the ureteric fibroblasts/smooth muscle) is derived from a distinct, Tbx18-positive progenitor, and thus might not be present in our isolated cells, we bioinformatically isolated interstitial cells from published datasets derived from whole kidneys (1,482 total cells) and included them in our analysis [17]. Unsupervised clustering was performed on all sequence d cells that met quality control standards. After pre-processing, quality control, normalization for cell cycle phase and lineage filtering, 8,683 interstitial cells were selected for further analysis.

Shared nearest neighbor clustering based on similarity between gene expression profiles revealed 17 distinct clusters of cells that were identified as “interstitial” (Figure 2, Supplemental Figure 1a, Supplemental Video 1 and Supplemental Video 2). Clusters ranged in size from 16 cells (cluster 13) to 1564 cells (cluster 10).

**Figure 2:**
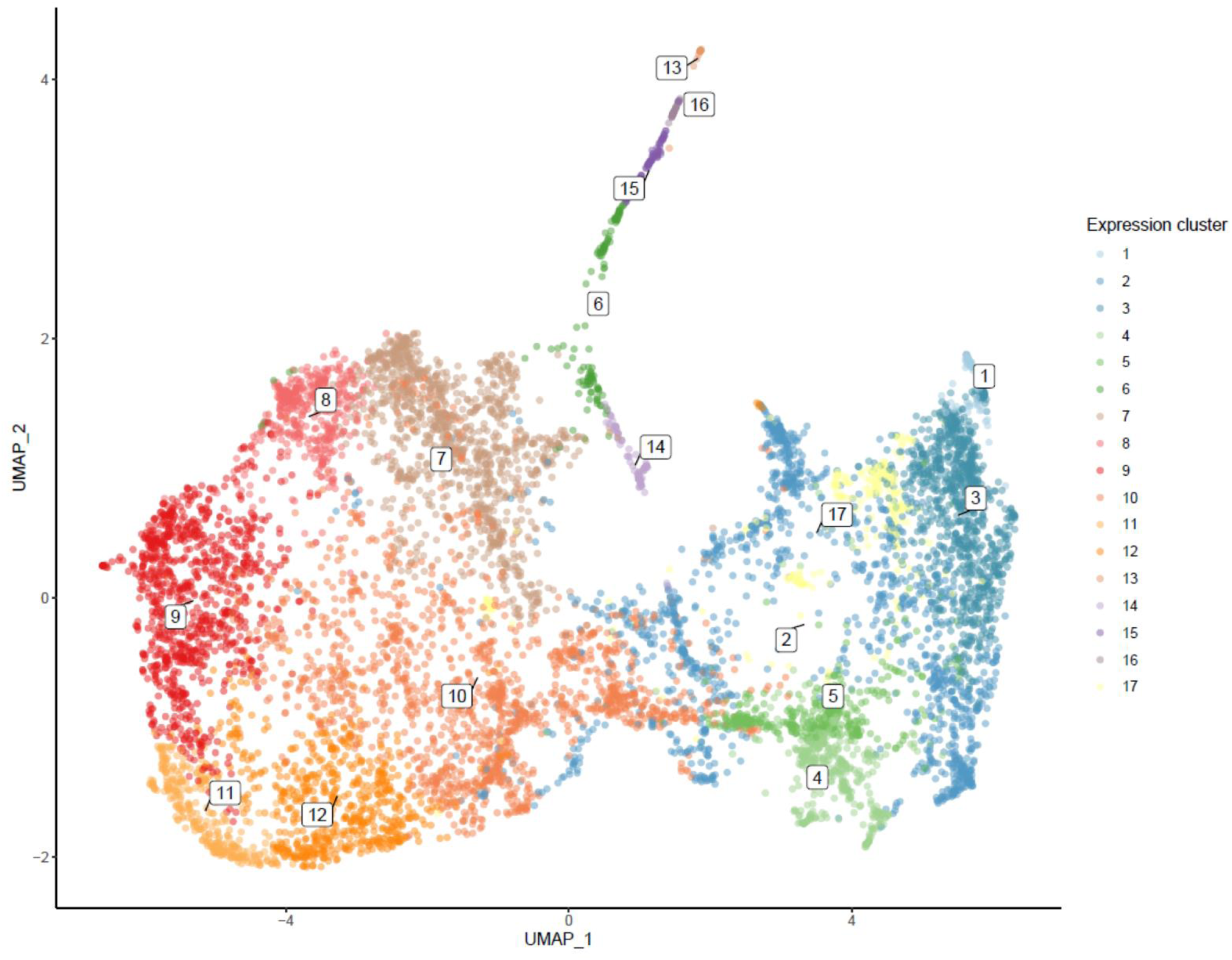
Interstitial heterogeneity revealed through single cell RNA sequencing: UMAP of E18.5 mouse intersitium. For a 3-dimensional representation of the UMAP see supplemental video 1 (a).

Importantly, the only cluster that was not derived from the Foxd1-Cre isolated cells was cluster 13, which represents the Tbx18-derived ureteric interstitium. These data suggest that our analysis includes most, if not all, renal interstitial cells, and that this population shows a high level of heterogeneity. We next sought to validate this heterogeneity in situ in order to gain insight into the significance and nature of the cellular diversity.

### Renal interstitium shows spatial heterogeneity

To validate our results, we performed section in situ hybridization with over 50 differentially expressed genes (DEGs) from each cluster (Figure 3-5, Supplemental Figure 1c-k and Supplemental Table 1, and Supplemental Table 2). Candidates were chosen based on relative abundance and any identified regionalized expression observed in publicly available databases [22–24]. Although all DEGs examined were expressed in the interstitium, some that were indicated as being differentially expressed between clusters did not show cell type specific expression by in situ hybridization (Supplemental Table 1 and Supplemental Table 2). Instead, these genes appeared to be ubiquitously expressed suggesting that their being called as a DEG was due to differences in mRNA levels between different cell types rather than cell type specificity. A subset of the queried genes were expressed broadly in epithelial and/or endothelial populations as well as the interstitium. Both of these classes of genes were largely excluded from further analysis. Several DEGs appeared to be enriched in the interstitium over other cell types and showed regionalized expression (see below). It is important to note that very few DEGs were detected in only a single cluster. Although individual examples for each cluster are presented in the main figures, anatomical assignment of clusters is based on in situ data from multiple DEGs (see supplemental Table 2). All in situ data will be available at the Re-Building a Kidney website (https://www.rebuildingakidney.org/).

**Figure 3:**
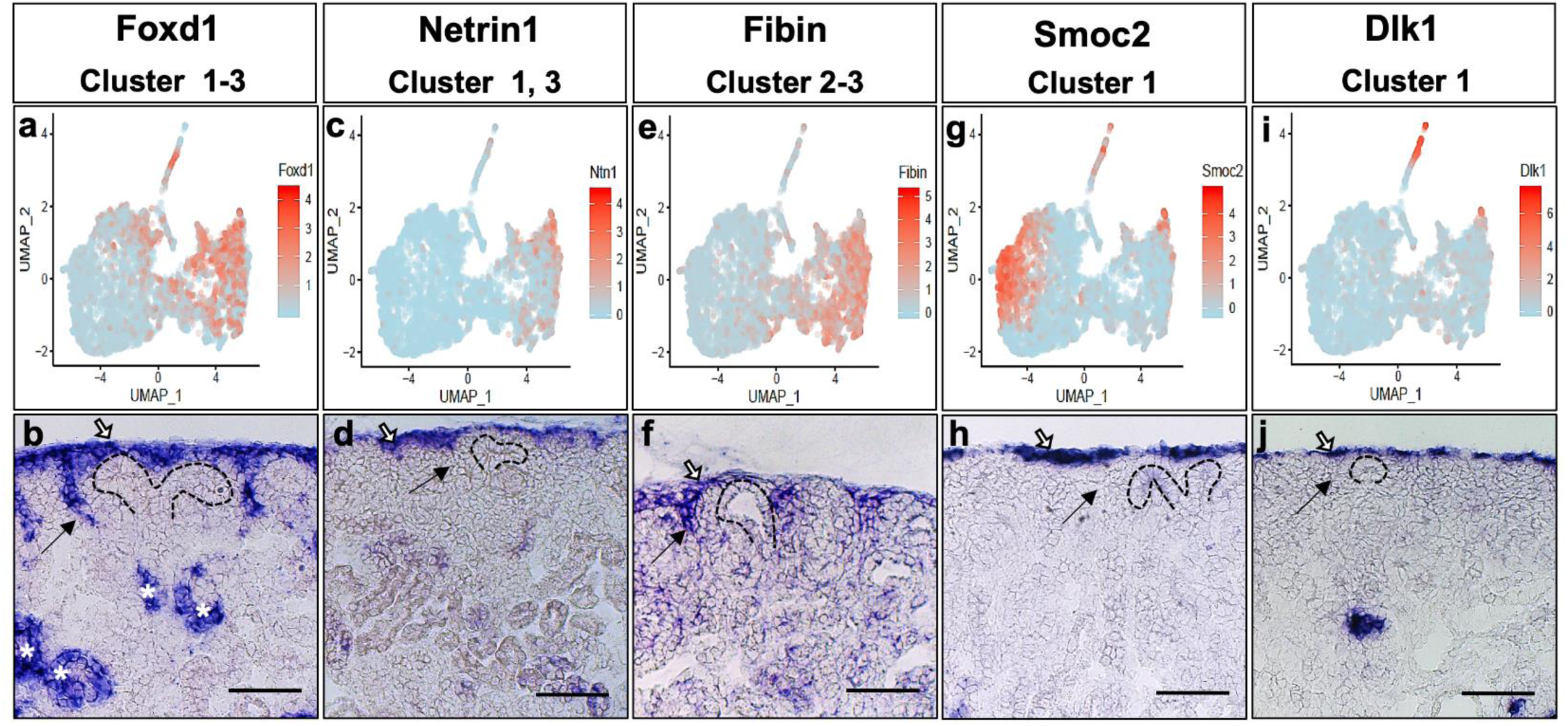
Single cell RNA sequencing reveals unexpected cortical heterogeneity within the renal interstitium that is validated through mRNA in situ hybridization. Specific gene expression displayed as a UMAP (a, c, e, g, i) for Foxd1(a), Netrin (c), Fibin (e), Smoc2 (g) and Dlk1 (i). E18.5 *in situ* hybridization of Foxd1, expressed in clusters 1-3 (b), Netrin1, expressed in clusters 1, 3 (d), Fibin expressed in clusters 2-3 (f), Smoc2, expressed in clusters 1 (h) and Dlk1, expressed in cluster 1 (j) reveals cortical heterogeneity. Scale bar: 100um

**Figure 4:**
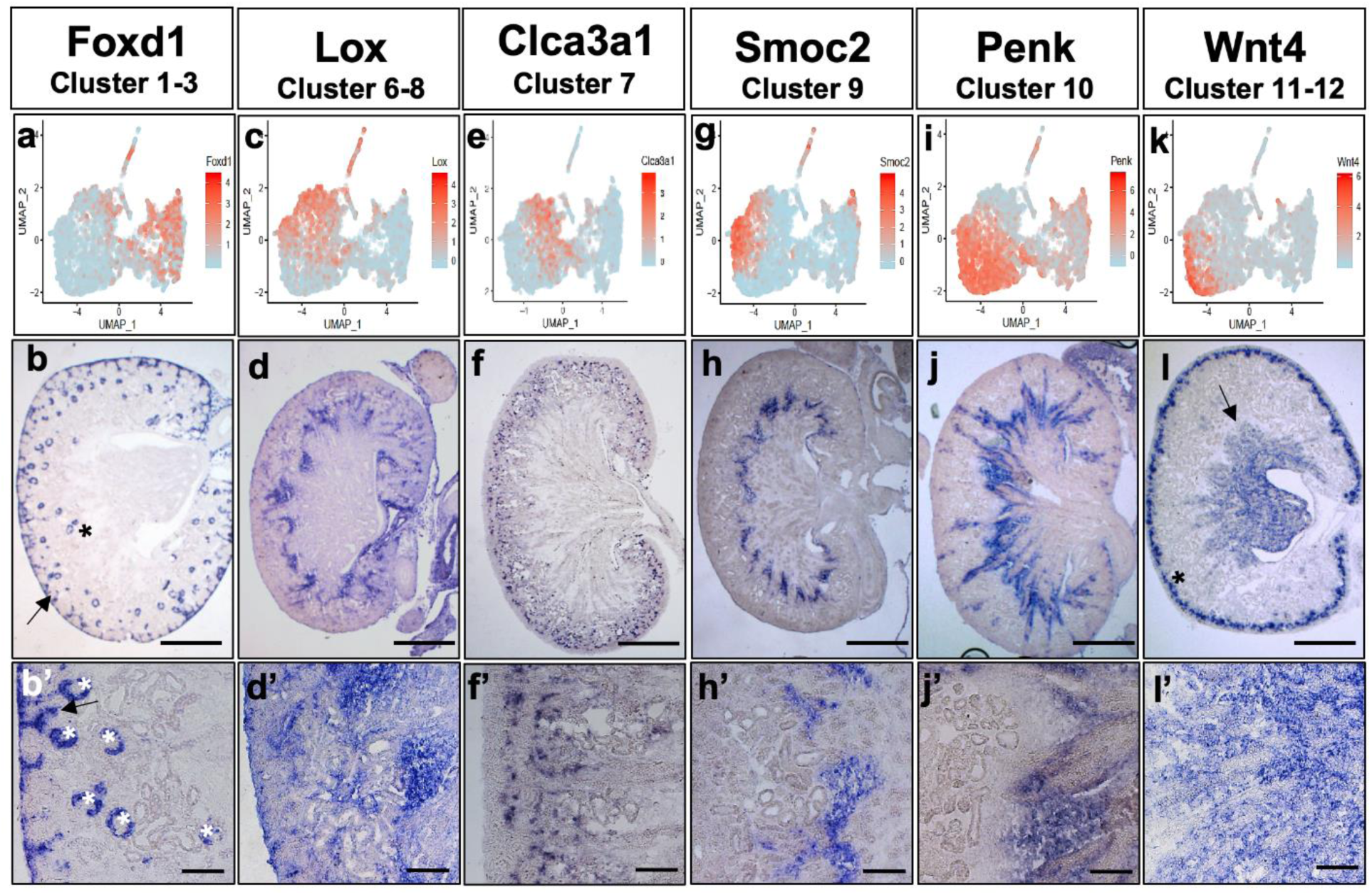
Spatial characterization of scRNA seq-clusters reveals cortico-medullary patterning among interstitial subtypes. Specific gene expression displayed as a UMAP (a, c, e, g, i, k) for Foxd1 (a), Lox (c), Clca3a1 (e), Smoc2 (g), Penk (i), and Wnt4 (k). E18.5 *in situ* hybridization of Foxd1: clusters 1-3 interstitial expression indicated with arrow, epithelial lineage expression in podocytes indicated with asterisk) (b-b’), Lox: cluster 7 (d-d’), Clca3a1: clusters 6-8 (f-f’), Smoc2: cluster 9 (h-h’), Penk: cluster 9-12 (j-j’), Wnt4: cluster 11-12 (interstitial expression indicated with arrow, epithelial lineage expression in PTA indicated with asterisk) (l-l’), reveals a molecular pattern which spans the cortico-medullary axis. b, d, f, h, j, l Scale bar: 500um. b’, d’, f’, h’, j’, l’ Scale bar: 100um.

**Figure 5:**
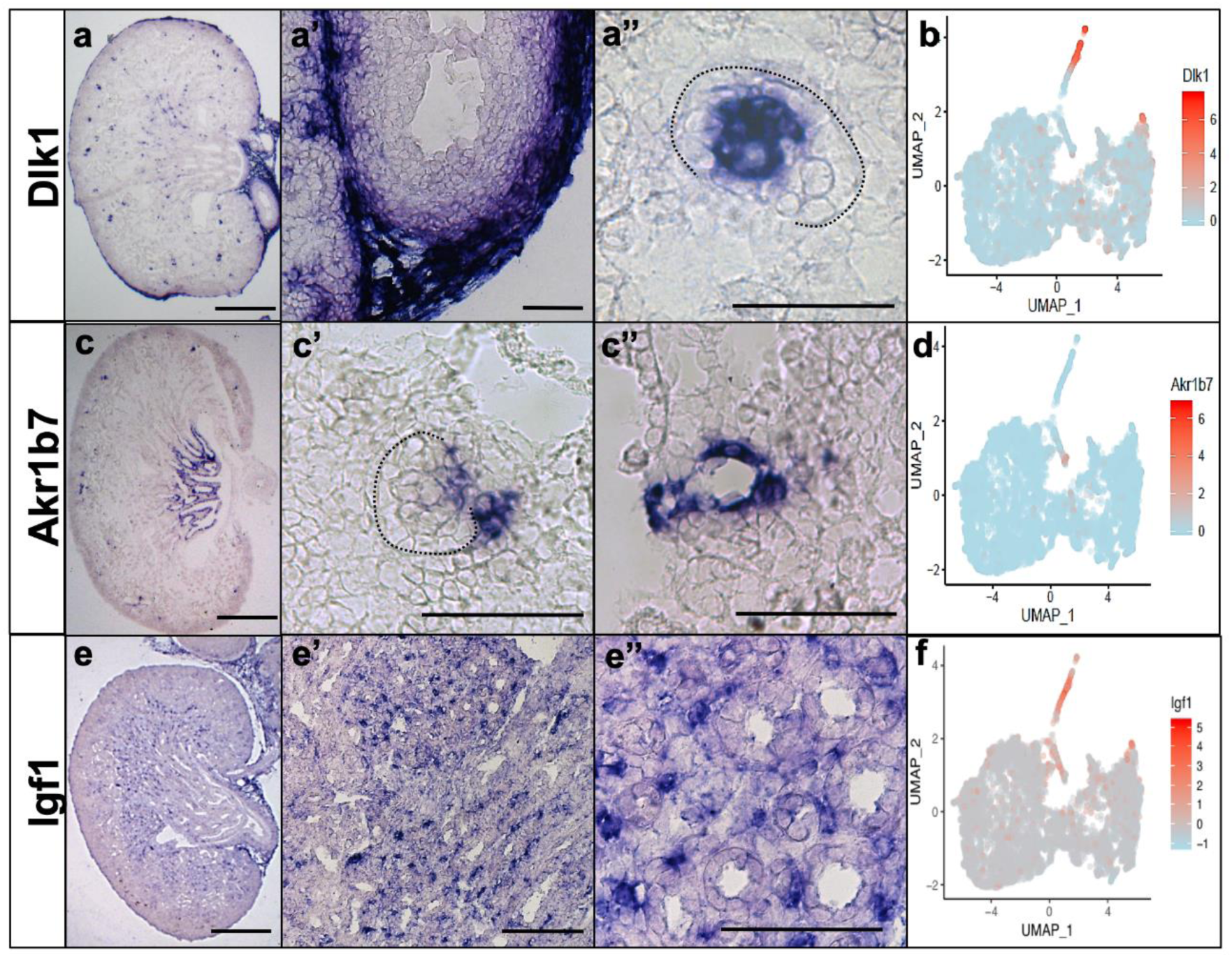
Mesangium and pericyte populations detected using single cell RNA sequencing. mRNA *in situ* hybridization of Dlk1 expression in clusters 15-16 (a) and Akr1b7 expression in cluster 14 (c). Dotted line outlines the glomerulus. Scale bar:100um Specific gene expression displayed as a tSNE (b, d) for Dlk1 (b) and Akr1b7 (d).

Clusters 1-3 are highlighted by expression of Foxd1, a gene known to be expressed in the cortical interstitium (Figure 4a-b, Supplemental Figure 1d, Supplemental Table 1) and podocytes, a non-interstitial cell type. Foxd1 expressing cells that were disjoint from interstitial cells and expressed typical podocyte markers, e.g. Podxl and Nphs2, were excluded from further analysis.

To gain insight into the nature of clusters 1-3, we assayed the expression of several mRNAs that were differentially expressed between the clusters including Netrin1 (clusters 1 and 3), Fibin (clusters 2 and 3), Smoc2 (cluster 1) and Dlk1 (cluster 1) (Figure 3, Figure 4a-b’, Supplemental Figure 1c-g, Supplemental Table 1). Netrin (clusters 1 and 3) was uniformly expressed in the interstitial cells cortical but not lateral to the cap mesenchyme (Figure 3c-d). Fibin (clusters 2 and 3), shows expression in a subset of the cortical interstitium as well as expression in cells that lie lateral to the cap mesenchyme (Figure 3e-f). Smoc2 (clusters 1 and 9) and Dlk1 (cluster 1, 13 and 16) both showed mosaic expression in the cortical most interstitium (Figure 3g-j) as well as other, non-cortical cell types (described below). Due to the limited resolution of in situ hybridization, we could not determine whether Fibin was expressed in distinct cell types from Smoc2 and Dlk1. However, this analysis suggests that the cortical Foxd1 expressing cells are not molecularly homogeneous and these observations suggest that clusters 1 and 3 are unique subsets within the cortical-most subset of Foxd1-expressing cells and cluster 2 represents a lateral sub-population of Foxd1-expressing cells. Thus, the region of stroma previously annotated as “cortical” appears to have at least 3 molecularly distinct cell types.

Genes present in clusters 4-5 were expressed on the medullary side of the ureteric bud tips surrounding the newly forming renal vesicles and correlated with a region of interstitium previously referred to as the nephrogenic interstitium (Supplemental Table 1 and data not shown). These clusters were highlighted by the expression of multiple genes involved in cell division (even after controlling for cell cycle phase).

Lysyl oxidase (LOX, clusters 6-8) was enriched in the interstitium adjacent to the most-medullary proximal tubules (Figure 4c-d’, Supplemental Figure 1h Supplemental Table 1). Clca3a1 (cluster 7) was also enriched in cells adjacent to the proximal tubules although it appeared not to be expressed adjacent to the more medullary proximal tubules and its expression appeared more mosaic than Lox (Figure 4e-f’).

Smoc2 (Cluster 9) is enriched in a population of interstitium that lies just medullary to the proximal tubule in the outer medulla (Figure 4g-h’, Supplemental Figure1g, Supplemental Table 1). Smoc2 expressing cells do not appear to expand cortically into the region adjacent to the proximal tubules.

The mRNA for Proenkephalin (*Penk*, clusters 9-12) is most intense in a population of interstitium just medullary to Smoc2 in the outer stripe of the inner medulla (Figure 4i-j’, Supplemental Figure 1i, Supplemental Table 1). Although its expression does appear to expand into more cortical populations of cells, it does not appear to overlap with Smoc2, Lox or Clca3a1.

Wnt4 is enriched in clusters 11-12. In situ analysis shows that along with expression in the pre-tubular aggregates, it is highly expressed in the interstitium of the papillary region of the kidney, a region previously referred to as the medullary interstitium. (Figure 4k-l’, Supplemental Figure 1j, Supplemental Table 1 and data not shown).

Cluster 13 is the only population that was not derived from the Foxd1 lineage sorted cells. All cluster 13 cells were derived from the whole kidney single cell data generated by Combes and colleagues [17]. Of note, cluster 13 was composed of only 16 cells, most likely a reflection of the relatively small numbers of interstitial cells included in the whole organ studies. As expected, genes present in cluster 13 (e.g. Dlk1 and Tbx18) were predominantly expressed in the interstitium adjacent to the ureter (Figure 1m, Figure 5a-b, Supplemental Figure 1c, Supplemental Table 1).

Clusters 14-16 showed a high degree of overlap in gene expression. All three clusters expressed several genes that are accepted markers of perivascular cells. To determine if the clusters represented unique perivascular cell types, we performed in situ hybridization with cluster specific DEGs. Akr1b7 and Ren1 are both enriched in cluster 14 over other clusters and show expression in vascular smooth muscle as well as cells within the juxtaglomerular apparatus (Figure 5c-d, Supplemental Figure 1b, Supplemental Table 1, and data not shown). Dlk1 (clusters 15 and 16 along with clusters 1 and 13) was detectable in the glomerular mesangial cells (Figure 5a-b, Supplemental Figure 1c, Supplemental Table 1). Igf1 was enriched in cluster 15 and showed expression in a subset of interstitial cells in the papillary region of the kidney, similar to the anatomical location assigned to cluster 10. Based on these observations along with gene set enrichment analysis [28] based on automated text-mining of protein-cell type associations from the biomedical literature [29] (Supplemental Figure 2), we suggest that cluster 14 represent vascular smooth muscle, cluster 15 represents a papillary pericyte population and cluster 16 represents mesangial cells. However, given the molecular similarity between these three cell types, it is possible that greater sequencing depth/clustering will alter these annotations.

Cluster 17 contained many genes that were also DEGs in other clusters. Although we did identify a specific DEG list for cluster 17, the most differentially expressed genes encoded mitochondrial mRNAs and we have not been able to produce strong signal with any of these probes (Supplemental Table 1).

In all, analysis of the transcriptome of single interstitial cells generated 17 clusters of cells that have been assigned to at least 12 anatomically distinct cell types.

### Pseudo-time analysis suggests multiple distinct developmental trajectories within the interstitium

Although interstitial heterogeneity is not completely unexpected, the heterogeneity in the histologically indistinguishable cells spanning the cortical-medullary axis was surprising. A simple explanation of the identity of these cells is that they merely represent different stages in a linear developmental trajectory from a cortical-most progenitor to a more medullary, differentiated cell type. To gain insight into the lineage relationship of the different clusters, we employed the complementary approaches of RNA velocity [25] and pseudotemporal ordering [26]. Briefly, RNA velocity is determined by modeling the relationship between the unspliced and spliced forms of a transcript, with the reasonable assumption that a cell actively transcribing a gene will have a higher ratio of unspliced to spliced transcripts and so a higher “RNA velocity”. Conversely, cells that are not actively transcribing a gene (but still maintain its expression) will have a lower ratio of unspliced to spliced transcripts as the majority of mRNA will represent processed transcripts (lower RNA velocity). Using single genes as representative examples, this analysis indicates that Lef1 is actively transcribed in cluster 10 and stabilizes in clusters 11 and 12 suggesting that clusters 11 and 12 represent cells derived from cluster 10 (Supplemental Figure 3a-b). In comparison, Smoc2 is actively transcribed and processed in cells present within cluster 9 but not in any other clusters (other than the Foxd1+ progenitor cells), suggesting cluster 9 represents a terminal differentiation state (Supplemental Figure 3c-d). Extending this model to all genes and all cells within the purified interstitium, we created an RNA velocity field to predict transitions between clusters (Supplemental Figure 3e). It is important to note that the kidney capsule, which is derived from Foxd1-expressing cells [7], was removed from our kidneys prior to cell sorting and thus is not included in our analysis. Therefore, we cannot comment on the derivation of this cell type at this time. We also ultimately excluded the ureteral smooth muscle cells (cluster 13) from this analysis as they did not show a relationship with any of the other clusters, as expected from their independent origins [21].

Simulation of a reverse Markov process with RNA velocity-based transition probabilities (please refer to Materials and Methods for a more complete description) allowed us to identify a group of cells that likely represented the ontological parent of the analyzed interstitial cells (Figure 6a). After selecting one such parent or “root” cell, we were able to pseudotemporally order the cells. Annotating the diffusion map with pseudotime demonstrated that the parent/root population ramifies into multiple distinct branches that develop simultaneously rather than following a single linear differentiation trajectory (Supplemental figure 3f).

**Figure 6:**
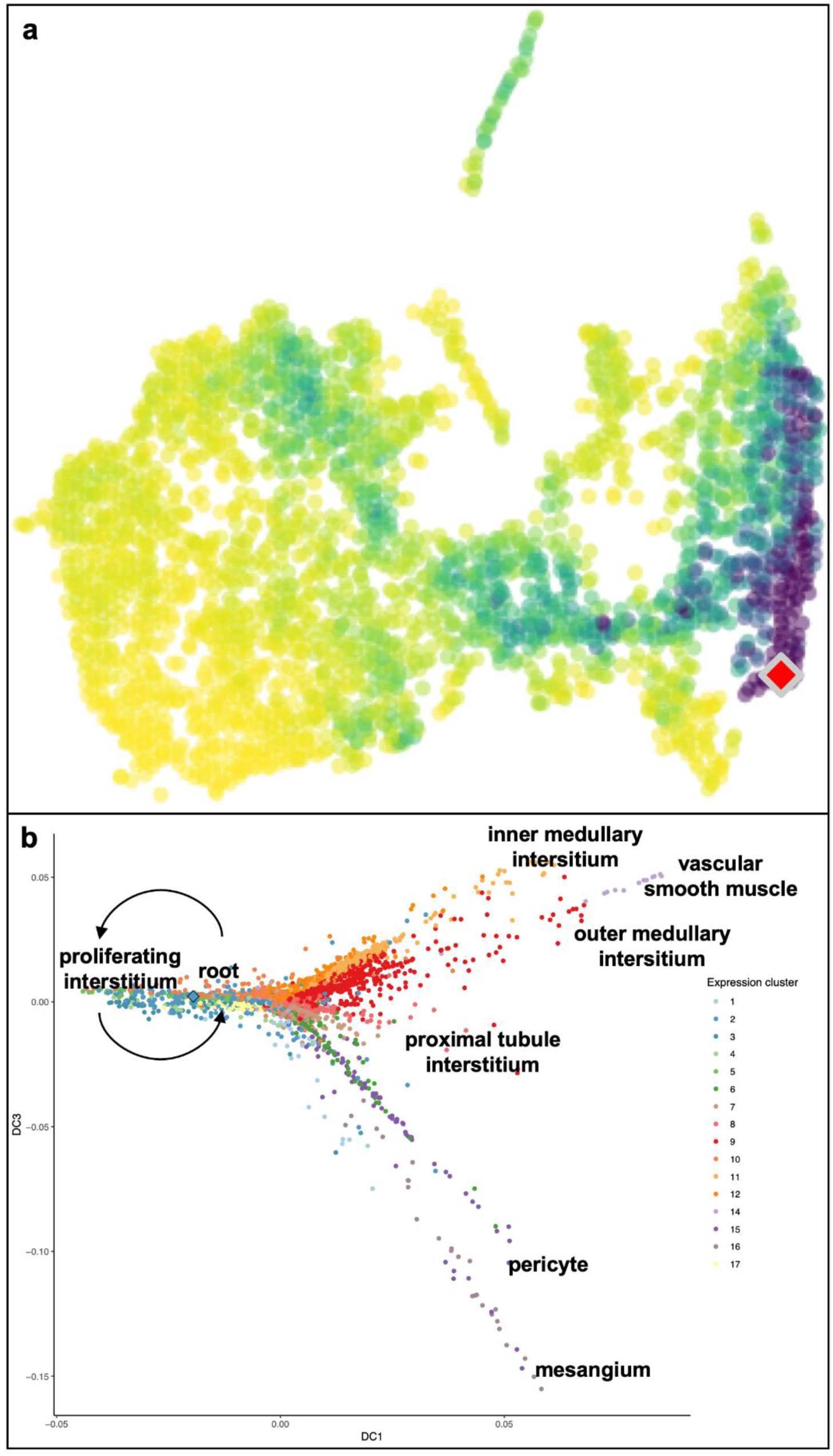
Foxd1 expressing cells may contain multiple, lineage-restricted progenitor cells. Differentiation roots identified by simulation of a Markov process with velocity-based transition probabilities in the reverse direction (a). Diffusion map of interstitial cells demonstrating dispersion of identified cell types along multiple distinct trajectories (b). For 3-dimensional representation of the diffusion map see supplemental video 3. The root cell is indicated by a diamond.

The integration of the RNA-velocity and pseudotime analyses generates a model wherein at least two of the distinct clusters of Foxd1-positive “progenitor” cells undergo independent branching events to give rise to the cycling/proliferating cells and at least 6 different trajectories that we refer to as the inner medullary fibroblast, outer medullary fibroblast, vascular smooth muscle, proximal tubule interstitium, pericyte and mesangial subtypes (Figure 6b). The vascular smooth muscle cluster appears more distantly related than the other cell types consistent with its distinct differentiation profile (Figure 6b and Supplemental video 3). Interestingly, clusters 6 appears to be located at a bifucation or trifucation point that gives rise to clusters 14, 15 and 16. This data suggests that the distinct interstitial subtypes do not represent transient stages in a linear developmental trajectory. Instead, these cells appear to represent previously undescribed, anatomically distinct interstitial cell types of distinct lineages.

Although frequently referred to as a homogeneous population of cells that functions primarily as a scaffold, several recent studies have shown that in various systems, the interstitium sets up unique microenvironmental niches that direct tissue development and/or maintenance and can contribute to pathological conditions [27–34]. Indeed, within the kidney, studies have shown that the interstitium is important in numerous developmental processes affecting both the epithelia and endothelia [5, 11–13]. We next sought to re-visit previous work in light of our current findings.

### Interstitial pattern affects development of the renal parenchyma

Several transcriptional regulators have been shown to be expressed in and necessary for the development of the stroma. To determine if any of these factors were active in specific stromal subtypes, we employed the SCENIC package to reconstruct gene regulatory networks and measure their activity within cells and so within the cells’ parent clusters (Figure 7) [35]. Strikingly, certain transcriptional signatures (aka regulons) were active in distinct clusters or groups of clusters while inactive in others. Interestingly, we found that the Lef1 regulon was predominantly active in clusters 8-12 (Figure 8a), which represents much of the medullary interstitium (Figure 5). Previous studies have shown that inactivation of beta-catenin in the interstitial progenitor population (using Foxd1Cre) leads to defects in the expression of several genes within the stroma, especially beta-catenin targets [36]. To test whether beta-catenin was necessary for formation of specific stromal subtypes (versus a general stromal defect), we assessed the expression of regionalized genes in Foxd1Cre;catnb^-/flox^ kidneys. Expression of genes normally expressed in clusters 1-7 (clusters where beta-catenin is not active) appeared unaffected or even expanded in Foxd1Cre;catnb^-/flox^ mutants (Figure 8b-d, h-j). In contrast, genes expressed in clusters 8-12 were markedly reduced or undetectable (Figure 8e-f, k-l) in mutants. The absence of medullary interstitium (clusters 8-12) correlated with a severe deficit in the formation of the epithelia (including the loop of Henle) that lay adjacent to these zones while the epithelia that lay adjacent to the unaffected cortical interstitium (including the proximal tubules) appeared to form normally, as previously reported by Yu et al. [36] (Figure 8g, m).

**Figure 7:**
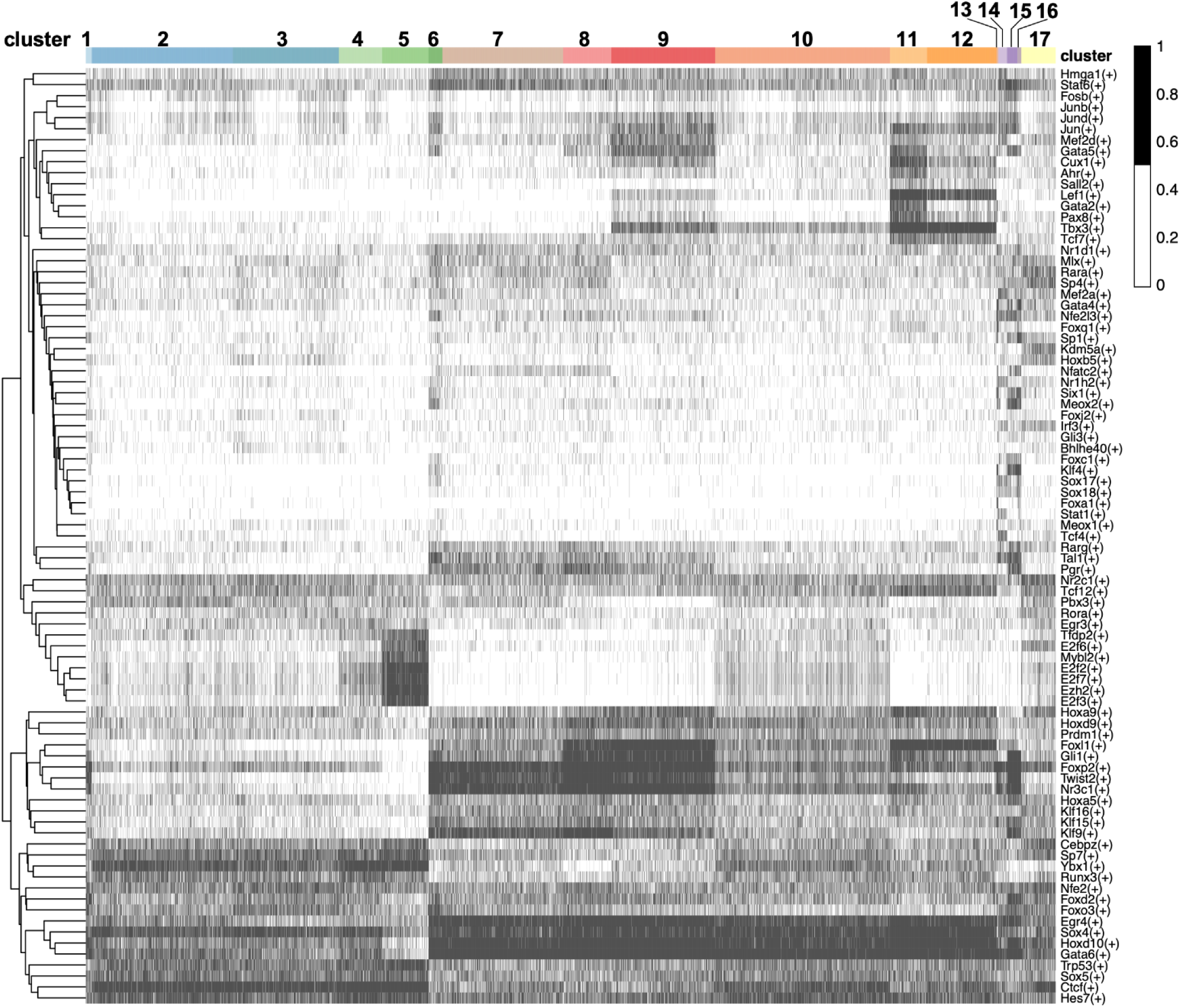
Distinct clusters show unique transcriptional activity. Heat map showing each cell organized by interstitial cluster (abscissa) and binarized regulon activity (ordinate) where black indicates an active regulon, while white indicates an inactive regulon.

**Figure 8:**
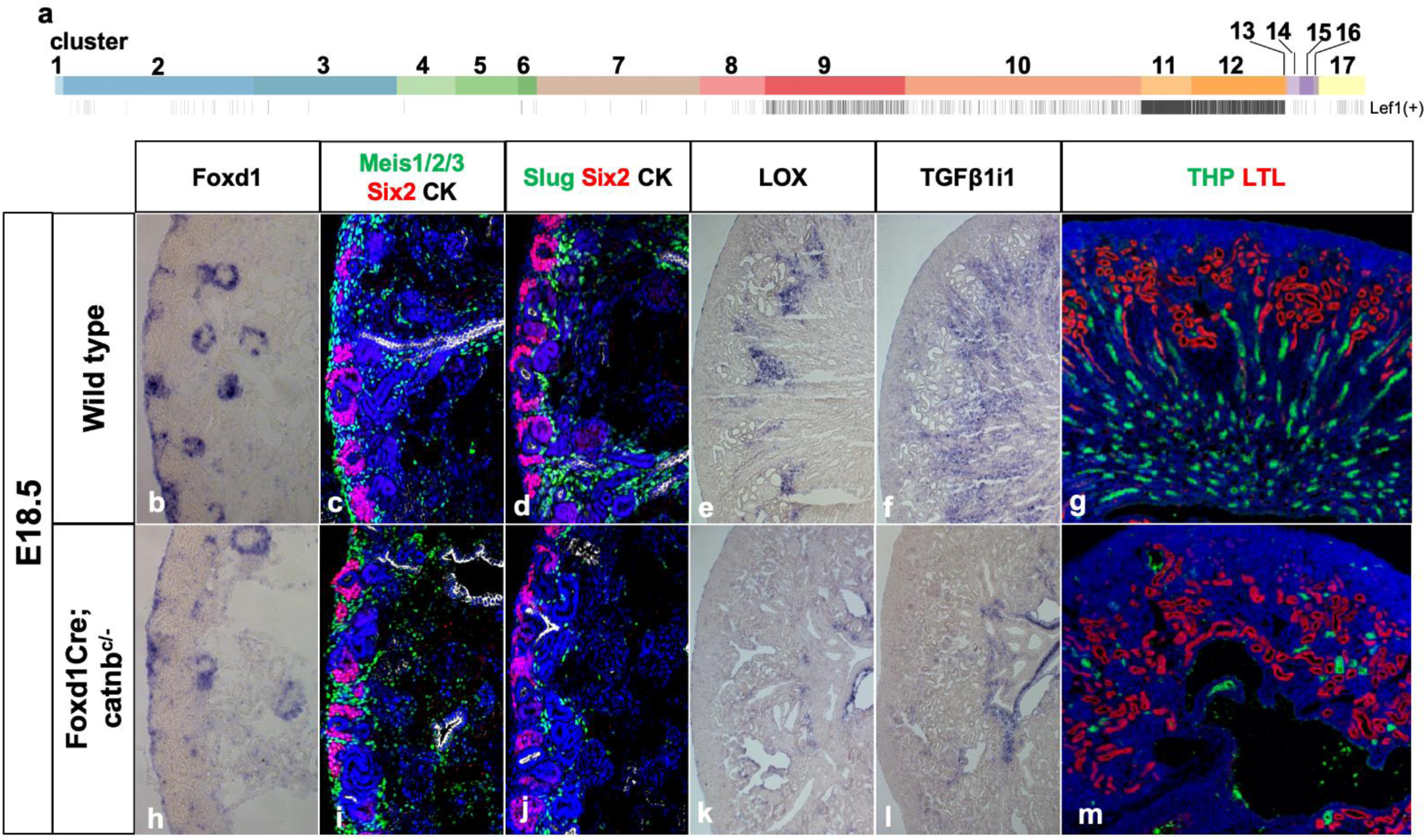
Perturbed patterning of the interstitial zones disrupts nephron differentiation. The Lef1 regulon is most active in clusters 9-12 (a). mRNA *in situ* hybridization (b, e, f, h, k, l) or immunofluorescence staining (c, d, g, i, j, m) of E18.5 wildtype (b-g) and Foxd1Cre;Beta-Catenin^c/-^ mutant (h-m) kidneys with probes to regional interstitial markers Foxd1(b, h), LOX (e, k) and TGFB1i1 (f, l) or antibodies to interstitial marker Meis1/2/3 (green in c, i) and Slug (green in d and j), nephron progenitor marker Six2 (red in c, d, i, j), collecting duct marker CK (white in c, d, i, j) or loop of Henle marker THP (green in g, m), and proximal tubule marker LTL (red in g, m).

### Human fetal kidney interstitium shows a similar degree of heterogeneity to the mouse

Recently, several groups have employed single cell RNA sequencing on human fetal kidney at different stages [37]. While analyzing the nephrogenic region of fetal kidneys, which contains cells from multiple lineages, only 5 unique interstitial sub-clusters were identified [37]. Thus, we wondered whether extensive interstitial heterogeneity is unique to the mouse.

To understand whether interstitial heterogeneity is a generalizable phenomenon between these two species, we reanalyzed previously published [38, 39] week 17 human fetal kidney scRNA seq data. After identifying the major cell populations within the data (epithelia, endothelia, leukocytes, etc.), we bioinformatically isolated the cells defined as interstitium (see methods). We then deployed the same clustering technique used to cluster mouse interstitium on the interstitium of the human fetal kidney. We find that the interstitium of the cortical region of human fetal kidney segments into 13 molecularly distinct clusters (Figure 9a). Although the human data set was limited to more cortical cell populations, we were able to identify almost as many unique clusters as found in whole embryonic kidney. When the cluster assignments from E18.5 mouse interstitial cells were mapped onto the 17-week human interstitial cells, the majority were found to be represented in the human data although they did not resolve as clearly (Figure 9b). There are likely several factors underlying this imperfect alignment including incomplete sampling in the human data, divergence between developmental time scales and time points between mouse and human and fewer cells available for analysis in the human dataset resulting in less resolving power. Evidence of the sampling bias is evident in that we did not identify any human cells that were analogous to cluster 13, the most medullary interstitium. These data indicate that renal interstitial heterogeneity is a generalizable characteristic between mouse and human.

**Figure 9:**
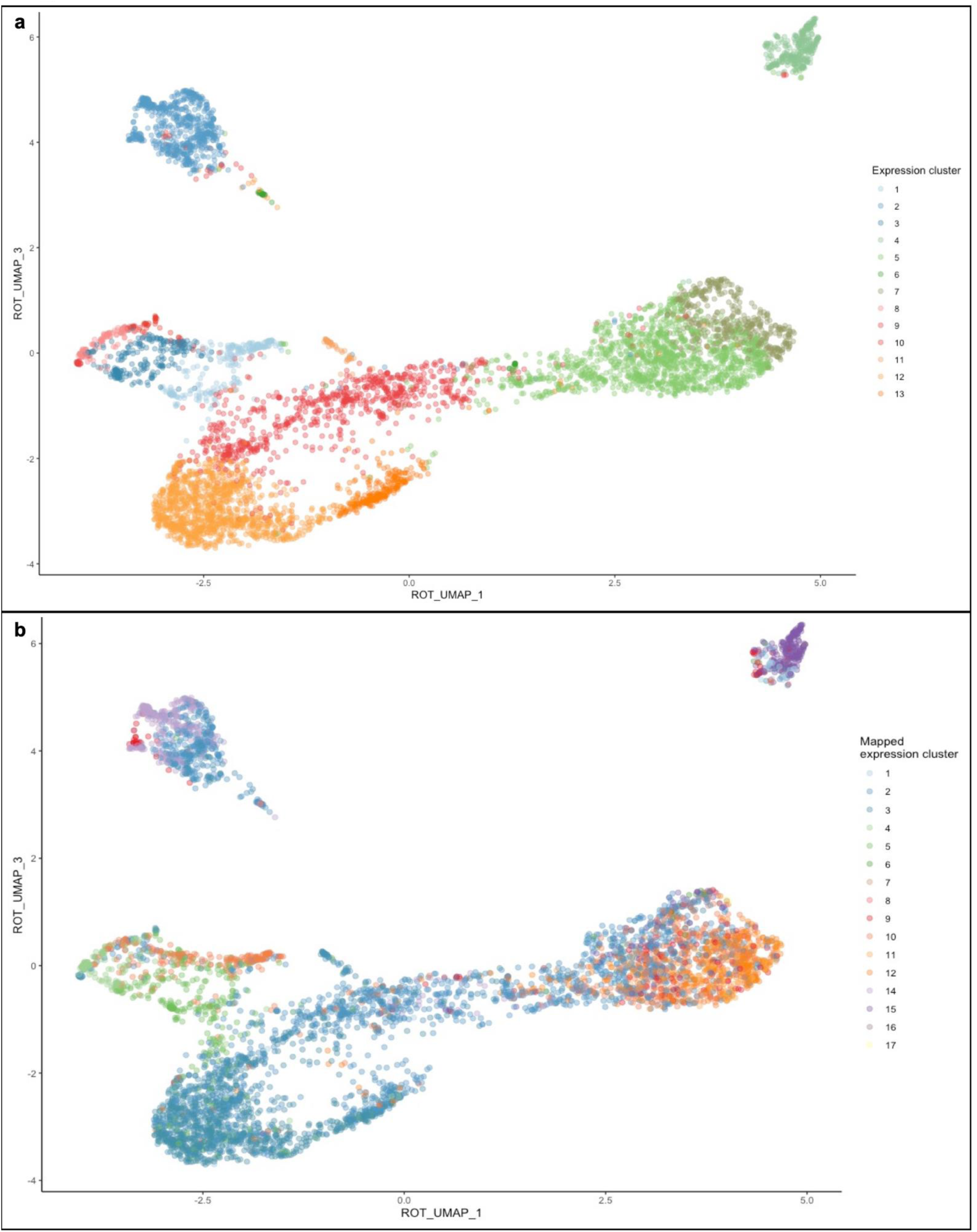
Human fetal renal interstitial heterogeneity revealed through single cell RNA sequencing: UMAP of 17 week fetal human renal intersitium. For a 3-dimensional representation of the UMAP see supplemental video 4 (a). UMAP of cluster assignments from E18.5 mouse interstitial cells mapped onto the 17-week human interstitial cells (b).

## DISCUSSION

Although known to play a role in providing physical support, growing evidence from multiple systems indicates that interstitial cells play active roles in tissue development, maintenance and disease. Within the developing kidney, non-autonomous roles for the interstitium have been identified in ureteric bud branching, nephron differentiation and blood vessel formation [5, 11–13]. Mechanistic insight into how the interstitium has so many distinct functions has been hampered by a poorly defined transcriptome. Here, using single cell RNA-sequencing combined with *in situ* hybridization, we have generated a map of interstitial gene expression in the E18.5 kidney. Our analysis shows a previously unappreciated level of heterogeneity in the interstitium of the developing mouse kidney. Analysis of human embryonic interstitial cells show correspondence in heterogeneity found in the embryonic mouse.

While previous work characterizing the heterogeneity of the developing mouse kidney uncovered 4 interstitial clusters (1,482 cells) [17], our informed analysis identified 17 distinct clusters that we were able to spatially resolve into at least 12 anatomically distinct subtypes. Additional analysis will be required to determine whether the 5 remaining clusters represent additional unique cell types or the data is currently overclustered. For example, we were not able to obtain signal from in situ hybridization with any DEGs specific to cluster 17, a population enriched for mitochondrial genes. Thus, it is possible that this cluster represents an artifact of the dissociation protocol. However, it is also worth noting that although immunostaining identified an interstitial cell type associated with the collecting ducts, our clustering did not conclusively identify these cells. It is possible that they lie within one of the medullary populations (12 or 13) or alternatively, that rather than being over-clustered, our data is under-clustered. We think it is likely that higher resolution techniques (e.g. single cell resolution in situ hubrization, antibody staining with or reporter gene generation from cluster specific DEGs) will resolve additional unique cell types. For example, there are three clusters containing genes expressed in the interstitium surrounding the proximal tubules. Although there do appear to be spatial differences in the expression of some proximal tubule interstitial genes, without cellular resolution, we cannot at this point determine whether these cells represent unique cell types. The fact that one of the, cluster 6, shows similarity to and is predicted to be the parent of clusters 14, 15 and 16, suggests that this cell type may represent a spatially distinct mural cell.

Unexpectedly, we found at least 3 molecularly distinct clusters within the Foxd1 expression domain. This observation raises the question of the identity of the true interstitial progenitor cell. One possibility that is supported by our RNA velocity analysis, is that rather than containing a single multipotent progenitor cell, the Foxd1 domain is comprised of several lineage restricted progenitors. Testing this will require more detailed molecular characterization as well as lineage tracing with cell type specific Cre drivers.

Interestingly, our analysis identified several zones of histologically indistinguishable but molecularly distinct fibroblast-like cell types that occupy unique spatial locations along the cortical medullary axis, where they correlate with distinct anatomical regions in the adjacent parenchyma. By analyzing our transcriptomic data, we were able to identify signaling pathways unique to the distinct interstitial clusters. Reanalysis of beta-catenin mutant interstitium reveals a unique role for this factor in the development of the papillary stroma, which secondarily affected the development of the adjacent epithelia. Of note, previous work has shown that mesenchymal cells play instructive roles in patterning the adjacent epithelia during development of various organ systems including patterning of the vertebrate gut tube [40–43]. The close correlation of the distinct kidney interstitial cell types with the functional subdomains within the nephron raises the intriguing possibility of stromal-epithelial cross talk that is involved in the patterning and/or differentiation of the kidney parenchyma. Although several studies have revealed cell autonomous mechanisms underlying nephron patterning, it is possible that distinct interstitial subpopulations produce factors that interact with intrinsic pathways to assure proper position and relative size and spacing of the nephron segments with the adjacent collecting ducts and associated renal vasculature. RNA-seq data reveals multiple growth factors, small molecules, extracellular matrix components and metabolites that appear to be regionally produced. Further genetic analysis will be required to test specific roles.

Finally, as the adult kidney shows exquisite patterning along the cortical/medullary axis, it will be of great interest to determine whether a similar degree of heterogeneity and pattern exists in the adult interstitium and how this pattern correlates to normal anatomy, physiology, injury, regeneration and disease state. Given the growing evidence of the essential nature of the interstitium in multiple processes [27–34], a similar analysis of interstitial heterogeneity in different organ systems at different stages may reveal that the interstitium’s role in patterning and morphogenesis is a generalizable principle.

## MATERIALS and METHODS

### Mice

All animals were housed, maintained and used according to protocols approved by the Institutional Animal Care and Use Committees at the University of Texas Southwestern Medical Center and following the guidelines from the NCI-Frederick Animal Care and Use Committee. For each experiment, female mice of 7–8 weeks of age were crossed with a male of 9–10 weeks of age. Plugs were checked and the embryos were collected at the desired time points for further analysis. Noon of the day on which the mating plug was observed was designated embryonic day (E) 0.5. The following mice were used in the studies described: Foxd1Cre (JAX Stock #012463), Rosa26Tomato (JAX Stock #007909), Rosa26DTA (JAX Stock #006331), Rosa26YFP (JAX Stock #006148), catnb null and catnb flox [44].

### In situ hybridization

For section *in situ* hybridization, kidneys isolated at specific stages were fixed overnight in 4% PFA (in PBS) at 4 °C and cryopreserved in 30% sucrose. Tissues were frozen in OCT (Tissue Tek) and sectioned at 10 μm. Sections were subjected to *in situ* hybridization as previously described [9]. The following antisense RNA probes against Foxd1, Lox, Smoc2, Penk, Wnt4, Lef1, Tgfb1i1, were linearized and transcribed as previously described. Plasmids were unavailable for Dlk1 and Akr1b7; thus, single stranded DNA for each gene was purchased with the T7 RNA polymerase binding site in the reverse orientation added to 3’ end of the gene sequence. These probes were made through RNA transcription of these single stranded DNA gblocks using T7 polymerase.

### Histology, immunohistochemistry and immunocytochemistry

Kidneys isolated at birth were formaldehyde fixed and paraffin embedded. Sections (5 μm) from paraffin-embedded kidneys were subjected to haematoxylin and eosin staining. For immunohistochemistry, fixed kidneys were embedded in OCT and sectioned on a cryostat (10 μm). Frozen sections were washed with PBS and blocked with 5% serum for an hour at room temperature and incubated with primary antibodies at 4 °C overnight. After primary incubation, sections were washed and incubated with HRP-tagged secondary antibodies for 1 h at room temperature. Further, signal was detected with tyramide amplification. Slides were washed and re-stained with additional markers according to the above-mentioned immunohistochemistry protocol. Slides were then mounted with Vectashield and images were captured with a Zeis LSM500, ZeisLSM700 or a Nikon A1R confocal microscope. The following antibodies were used:

| Protein Target | Antibody Product ID | Dilution |
| --- | --- | --- |
| Acta2 (aSMA) | Sigma-Aldrich Cat# A2547, RRID:AB_476701 | 1:250 |
| Cdkn1c | ThermoFisher Cat# PA5-34432, RRID:AB_2551784 | 1:250 |
| DBA | Vector Laboratories Cat# B-1035, RRID:AB_2314288 | 1:500 |
| E-Cadherin | Invitrogen Cat#13-1900, RRID:AB_2533005 | 1:250 |
| E-Cadherin | BD Biosciences Cat# 610182, RRID:AB_397581 | 1:250 |
| Foxd1 | Santa Cruz Cat# SC-47585; RRID:AB_2105295 | 1:750 |
| GFP | Aves Labs Cat# GFP-1020, RRID:AB_10000240 | 1:500 |
| Laminin | Sigma-Aldrich Cat# L9393, RRID:AB_477163 | 1:500 |
| Lef1 | Cell Signaling Technology Cat# 2230, RRID:AB_823558 | 1:250 |
| LTL | Vector Laboratories Cat# B-1325, RRID:AB_2336558 | 1:500 |
| Meis1/2 | Santa Cruz Cat# SC-10599, RRID:AB_2143020 | 1:750 |
| Meis1/2/3 | Active Motif Cat# 39795, RRID:AB_2750570 | 1:100 |
| pan Cytokeratin<br>(CK) | Sigma-Aldrich Cat# C2562, RRID:AB_476839 | 1:500 |
| PDGFrβ | Cell Signaling Cat# 3169S, RRID:AB_2162497 | 1:250 |
| Six2 | Proteintech Group Cat# 11562-1-AP, RRID:AB_2189084 | 1:500 |
| Slug | Cell Signaling Technology Cat# 9585, RRID:AB_2239535 | 1:250 |
| Tbx18 | Santa Cruz Cat# SC-17869, RRID:AB_2200374 | 1:100 |
| THP | Alfa Aesar Cat# J65429 | 1:500 |

### Single cell sample preparation and sequencing

E18.5 Foxd1Cre;Rosa26Tomato mouse kidneys were dissected in cold PBS without calcium or magnesium, Kidneys were cleaned and adrenal gland, capsule and ureters were removed. Kidneys were washed in HBSS for 2 minutes at 37°C then minced using two razorblades on ice. The minced kidneys were digested for 8 minutes at 37C in 2mL of 0.25% w/v Collagenase A/1% w/v Pancreatin (Sigma-Aldrich; Cat. 101378001, P1750), with manual dissociation via pipetting through a P1000 tip every 2 minutes. No more than 4 pairs of kidneys were dissociated in 2mL enzyme digest. Digestion was inactivated by adding 125ul serum. After pelleting the cells at 400g for 5 minutes, cells were resuspended in 1mL of AutoMACS Running Buffer (Miltenyi Biotec, Cat. 130-091-221) and passed through a 30 um pre-separation filter (Milteni Biotec, Cat. 130-041-407). Filters were immediately washed with 500ul AutoMACS Running Buffer. Cells were resuspended in 500uL AutoMACS Running Buffer and filtered at least 2 more times through a cell-strainer cap (Falcon, Cat. 352235) attached to a 5mL Polypropylene round bottom tube (Globe Scientific, Cat. 110428). Ten thousand Foxd1Cre;Rosa26Tomato+ cells, isolated via FACS, were run on a chromium 10x Single Cell Chip (10x Genomics). Libraries were prepared using Chromium Single Cell Library kit V2, and sequenced on an Illumina NextSeq using 75pb paired-end sequencing. Sequencing resulted in an average of 12,000 reads/cell and 3,000 genes/cell. Upon acceptance, the single cell data presented in this manuscript will be deposited onto Gene Expression Omnibus and Rebuilding a Kidney databases.

### Single cell data analysis

When analyzing the single cell data collected from the experiments outlined above, we included, where possible, the dataset generated by Combes et al. obtained via NCBI’s Gene Expression Omnibus under accession GSE108291 [17]. Each batch was processed independently using the scran Bioconductor package [45]. Unfiltered feature-barcode matrices were generating by running the CellRanger count pipeline. Cells were called from empty droplets by testing for deviation of the expression profile for each cell from the ambient RNA pool [46]. Cells with large mitochondrial proportions, i.e. more than 3 mean-absolute deviations away from the median, were removed. Cells were pre-clustered, a deconvolution method was applied to compute size factors for all cells [47] and normalized log-expression values were calculated. Variance was partitioned into technical and biological components by assuming technical noise was Poisson-distributed and attributing any estimated variance in excess of that accounted for by a fitted Poisson trend to biological variation. The dimensionality of the data set was reduced by performing principal component analysis and discarding the later principal components for which the variance explained was less than variance attributable to technical noise.

Masking of biological effects by expression changes due to cell cycle phase were mitigated by blocking on this covariate. The cell cycle phase was inferred using the pair-based classifier implemented in the *cyclone* function of scran. Corrected log-normalized expression counts were obtain by calling the removeBatchEffect from the limma [48] Bioconductor package with a design formula including G1 and G2M cell cycle phase scores as covariates.

A single set of features for batch correction were obtained by computing the average biological component of variation across batches and retaining those genes with a positive biological component. The batches were rescaled and log-normalized expression values recomputed after the size factors were adjusted for systemic differences in sequencing depth between batches. Batch effects were corrected by matching mutual nearest neighbors in the high-dimensional expression space [49]. The resulting reduced-dimensional representation of the data was used for all subsequent embeddings including t-SNE and UMAP.

Cells were clustered by building a shared nearest neighbor graph[50] and executing the Walktrap algorithm [51]. Differential gene expression analysis was performed using the two-part generalized linear model that concurrently models expression rate above background and expression mean implemented in MAST[52]. A one-versus-all strategy was employed comparing each cluster to all other identified interstitial clusters.

Gene sets for enrichment analysis were obtained from the TISSUES Text-mining Tissue Protein Expression Evidence Scores datase t[53] located at http://amp.pharm.mssm.edu/Harmonizome/. The gene sets were filtered to include only those genes with a standardized value greater than 1. Enrichment analysis was performed using the fgsea [54] Bioconductor package.

Week 17 human fetal kidney scRNA seq data was obtained from the Gene Expression Omnibus under series accession numbers GSE112570 and GSE124472 [38, 39]. Cluster assignments were transferred from the E18.5 mouse kidney dataset to the 17-week human fetal kidney dataset using a neural network classifier constructed using the Tensorflow system [55]. Graph regularization [56] considering eight shared nearest neighbors of each cell was used during training of a sequential network with two hidden layers, each containing 1024 hidden nodes, to classify the expression profiles of cells from the mouse dataset by cluster. Orthologue-mapped expression profiles of human cells as input to the classifier to assign each cell in the human dataset to mouse cluster with greatest similarity. Expression profiles were cosine-normalized prior to training or prediction. Only genes with a biological-to-technical variance ratio greater than zero were utilized for classification.

RNA velocity was calculated to analyze the dynamic relationships between identified cell states [25]. The analysis was limited to the replicate single cell datasets produced for this paper as BAM files were not available for the Combes, *et al*. dataset [17]. Read counts were partitioned between spliced, unspliced and ambiguous sets by calling velocyto *run10x* with an mm10 repeat mask obtained from the UCSC genome browser and the genome annotation file that came prepackaged with cellranger. The results of previously conducted cell filtering and feature selection were applied to these expression matrices. Normalization, principal component analysis, k-nearest neighbor smoothing, gamma fit, extrapolation, Markov process modelling and projection onto pre-defined low-dimensional embeddings were executed by calling the relevant functions from the velocyto.analysis module.

To reconcile clustering with trajectory inference, we performed partition-based graph abstraction (PAGA) to model the connectivity between clusters [57]. At cluster resolution, the edge-score threshold was varied until the graph was decomposed into connected components. Evaluation of marker gene expression within the components permitted assignment of components to the categories epithelium, leukocyte, erythrocyte, endothelium and interstitium. In many cases, RNA velocity allowed for assigning a direction to the edges between clusters indicating a tendency of transition between the clusters.

Gene regulatory network reconstruction and measurement of regulon activity within each was conducted using SCENIC [35]. All cells of acceptable quality were used for network inference. The regulon activity was binarized using the following strategy. An attempt was made to model the regulon’s activity as a mixture of two normal distributions using the mixtools R package. If such a model could be fit using expectation maximization, then those cells for which the regulon activity were assigned higher probability by the distribution with the greater mean were assigned a binarized regulon activity of one and zero otherwise. If a two-component mixture of normal distributions could not be fit, then a beta distribution was fit to the regulon’s activity and cells for which the regulon’s activity was greater than one mean absolute deviation above the mean were assigned a binarized regulon activity of one and zero otherwise.

## Acknowledgements

The authors would like to thank Ondine Cleaver, Denise Marciano, and Phoebe Carter for their insight in preparing this manuscript.

## Data Availability

Upon acceptance, the single cell data presented in this manuscript will be deposited onto Gene Expression Omnibus and Rebuilding a Kidney databases with DOIs included.

## Author Contributions

ARF designed experiments, performed experiments, analyzed data and wrote the manuscript. CPC designed experiments, analyzed data and wrote the manuscript. AD designed experiments, performed experiments and analyzed data, KD and MP designed experiments. CSK designed experiments, performed experiments, analyzed data and wrote the manuscript. AM, DA, DS, and GH prepared single cell libraries. TJC designed experiments, analyzed data and wrote the manuscript.

---

**Supplemental Figure 1:**
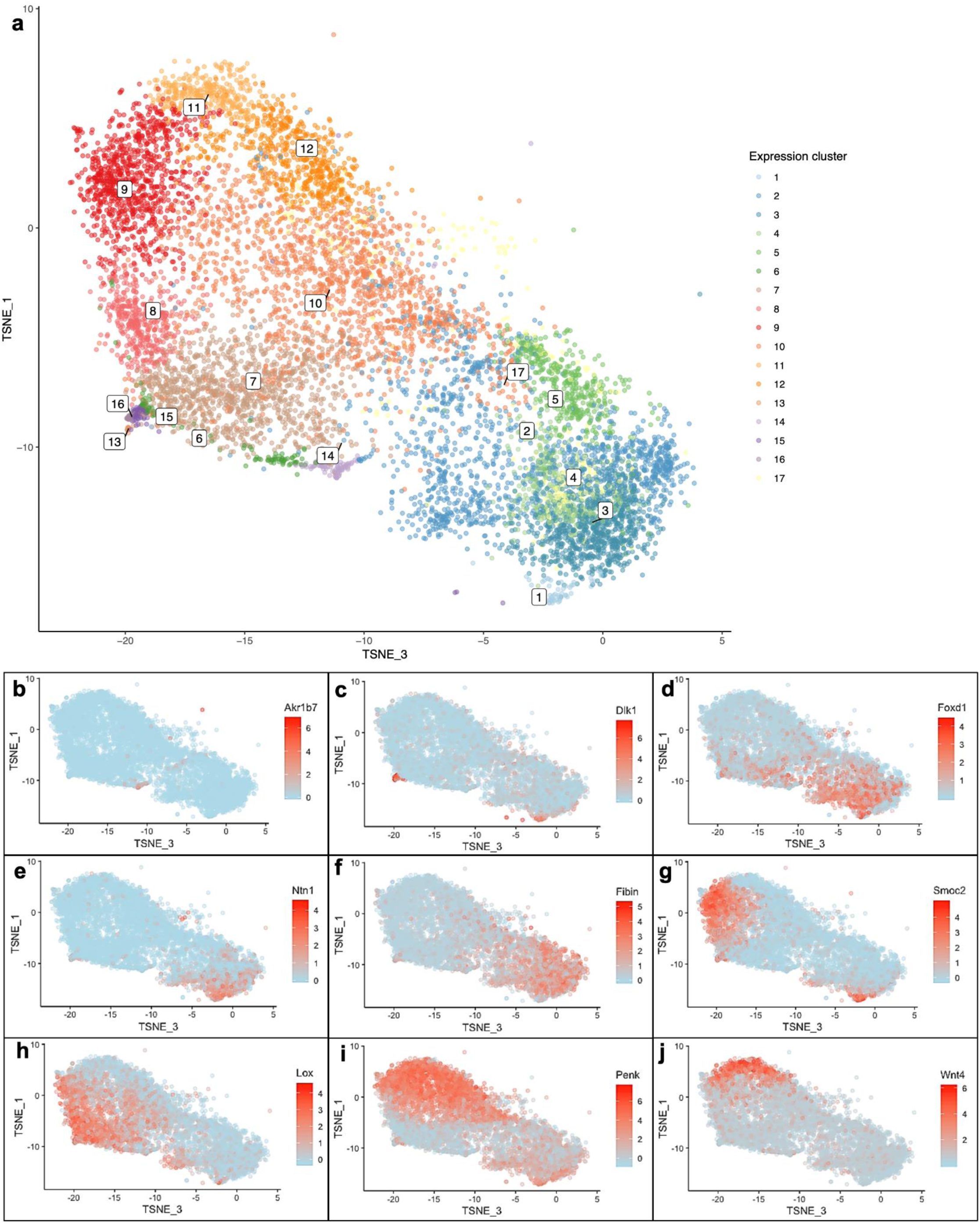
tSNE of renal interstitium. For a 3-dimensional representation of the tSNE see supplemental video 2 (a). Specific gene expression displayed as a tSNE (b-j) for Akr1b7 (b), Dlk1 (c), Foxd1 (d), Ntn1 (e), Fibin (f), Smoc2 (g), Lox (h), Penk (i) and Wnt4 (j)

**Supplemental Figure 2:**
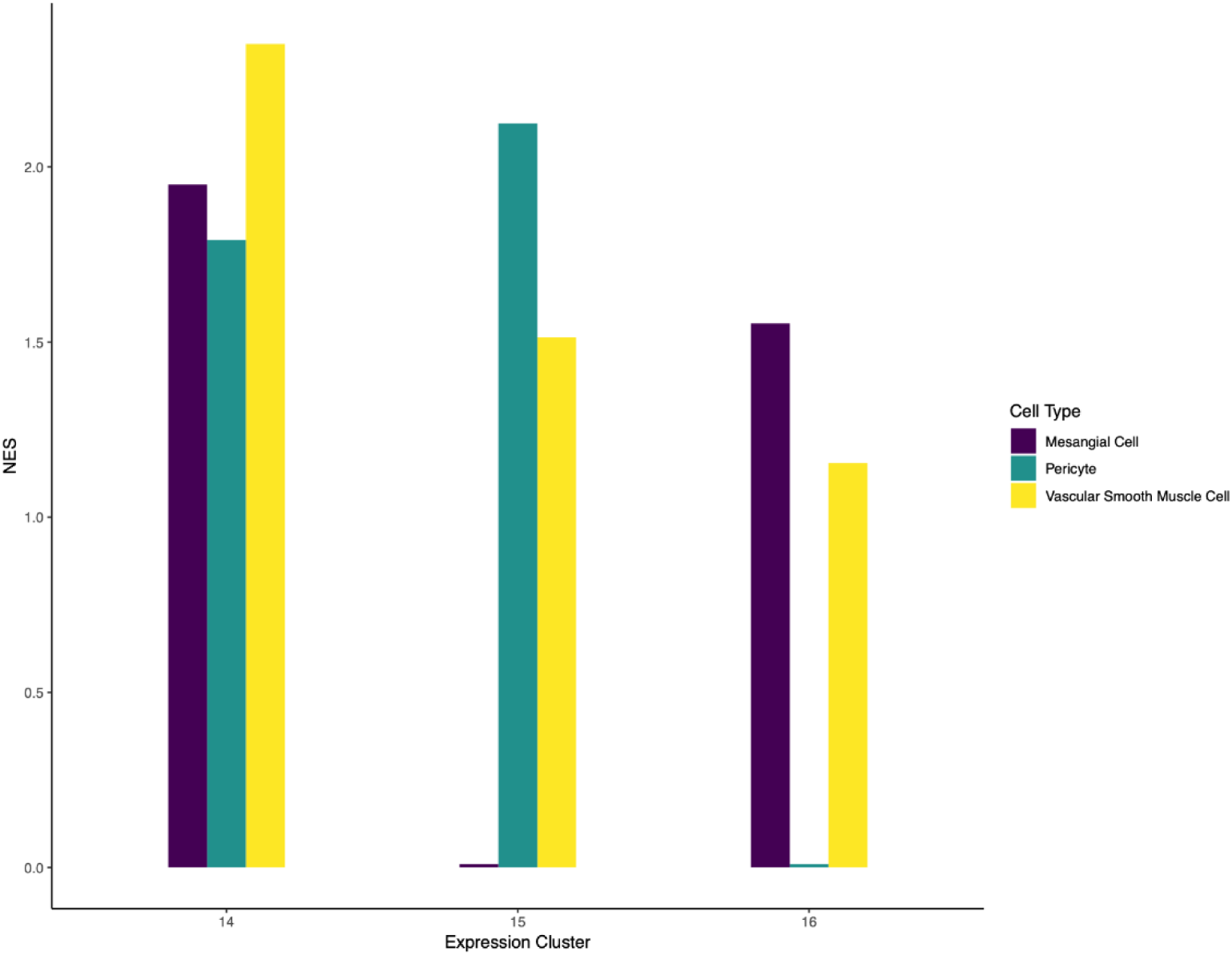
Normalized enrichment scores (NES) for mesangial, pericyte and vascular smooth muscle cell types specifically for clusters 14, 15 and 16.

**Supplemental Figure 3:**
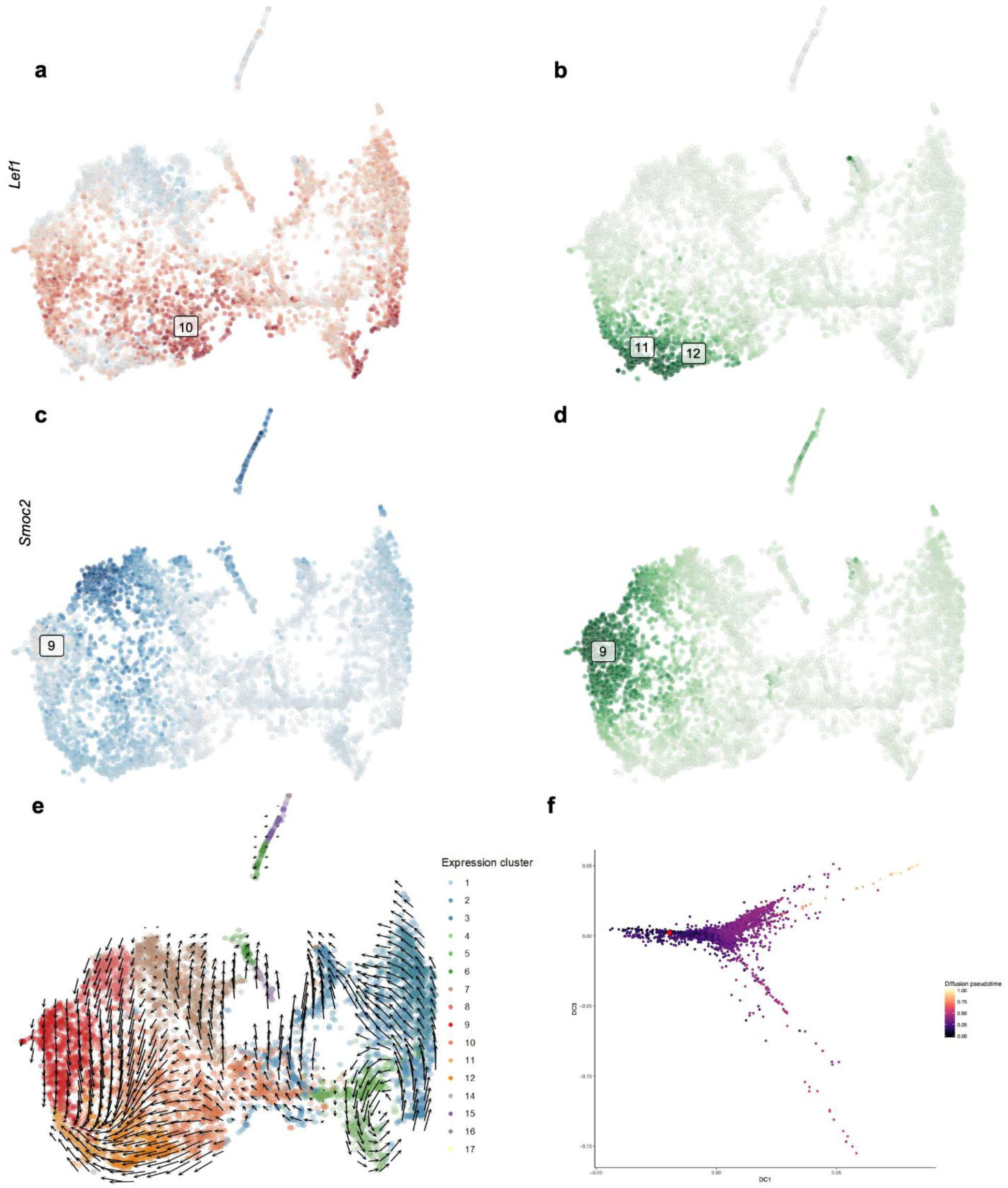
UMAP with cells colored by abundance of Lef1 (a) or Smoc2 (c) unspliced transcripts. UMAP with cells colored by abundance of Lef1 (b) or Smoc2 (d) spliced transcripts. RNA velocity field visualized using Gaussian smoothing on a regular grid (e). Diffusion map of stoma cells colored by diffusion pseudotime (f).

**Supplemental Table 1:**
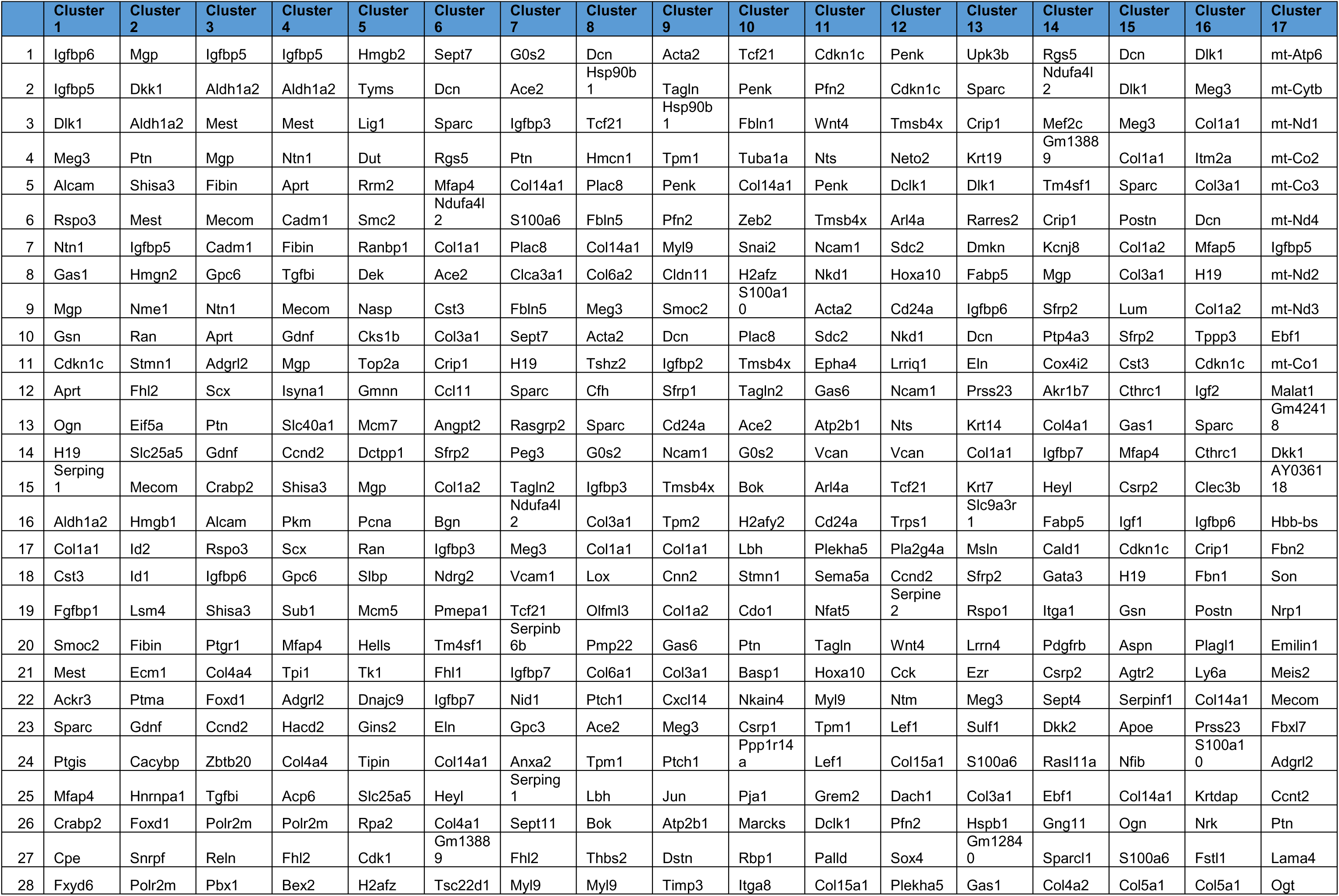
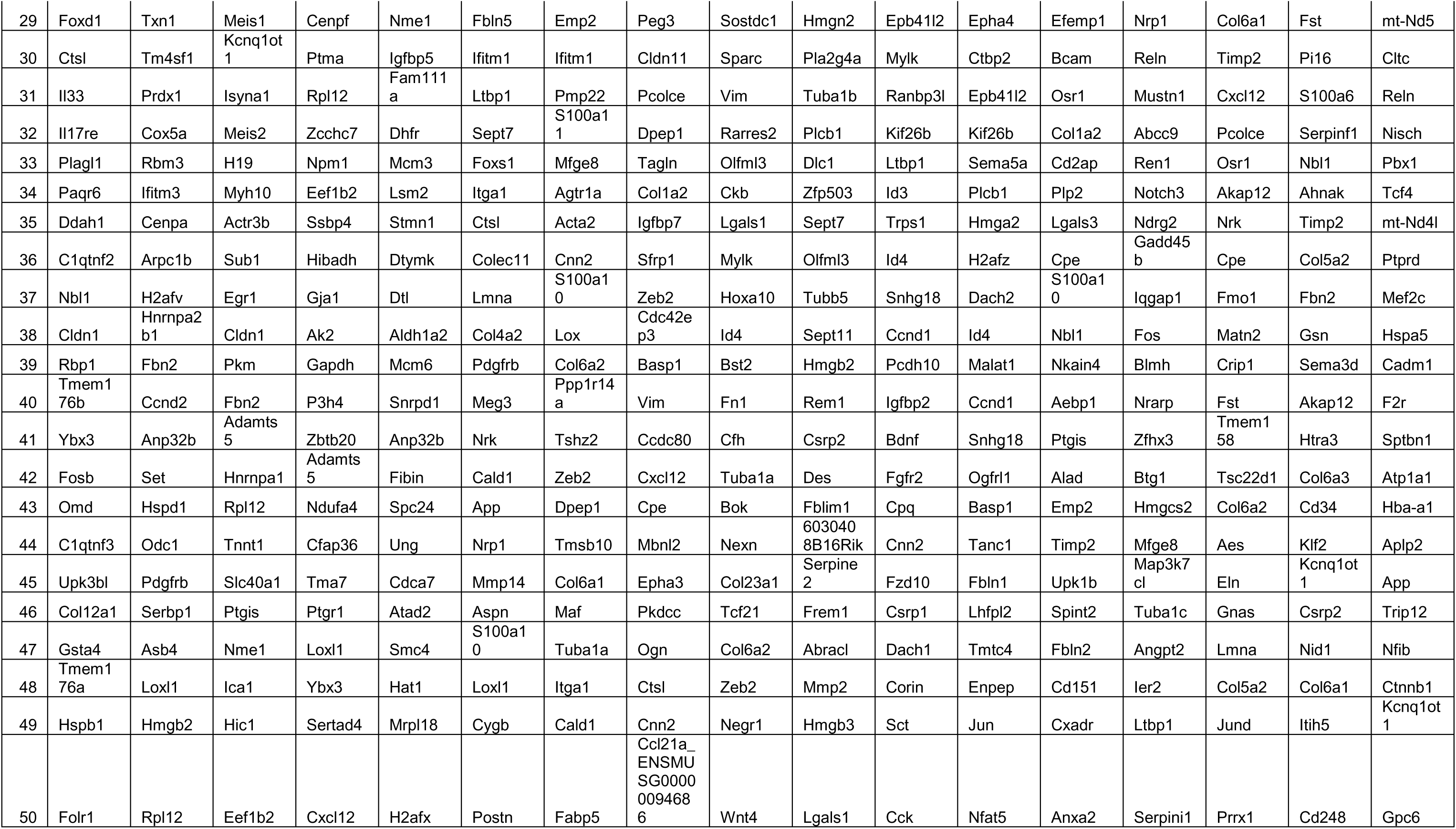
List of top 50 differentially expressed genes (DEGs) in each cluster when compared to all clusters. Our annotations for each cluster as described in the text are as follows: Cortical Interstitium: Clusters 1-3; Nephrogenic Interstitium: Clusters 4-5; Proximal Tubule Interstitium: Clusters 6-8; Interstitium Medullary to Proximal Tubule (Outer Medulla): Cluster 9; Outer strip of inner medulla Interstitium: Cluster 10; Papillary Interstitium: Clusters 10-12; Ureteric Interstitium: Cluster 13; Vascular Smooth Muscle: Cluster 14; Pericyte: Cluster 15; Mesangium: Cluster 16; Indeterminate Signature: Cluster 17.

**Supplemental Table 2:** List of stromal gene against which mRNA in situ hybridizations were conducted, the image of the in situ, along with the rank of the gene in the DEG list for each cluster. Yellow indicates the gene was ranked in the top 100 DEGs, while grey indicates the gene was not present in the DEG list for that specific cluster.

**Supplemental Video 1:** 3-dimensional view of mouse renal interstitial UMAP

**Supplemental Video 2:** 3-dimensional view of mouse renal interstitial tSNE

**Supplemental Video 3:** 3-dimensional view of mouse renal interstitial diffusion map

**Supplemental Video 4:** 3-dimernasional view of fetal human renal interstitial UMAP.

